# Efficient protein structure prediction from compact computers to datacenters with OpenFold-TRT

**DOI:** 10.64898/2026.03.11.711233

**Authors:** Kieran Didi, Prashant Sohani, Fabian Berressem, Alexander Nesterovskiy, Boris Fomitchev, Robert Ohannessian, Mohamed Elbalkini, Jonathan Cogan, Anthony Costa, Arash Vahdat, Felix Kallenborn, Bertil Schmidt, Milot Mirdita, Martin Steinegger, Christian Dallago, Alejandro Chacon

**Author notes:** Correspondence: {, }. Equal contribution (Kieran Didi, Prashant Sohani, Alejandro Chacon).

## Abstract

We introduce accelerations for deep learning inference with OpenFold and TensorRT that, combined with MMseqs2-GPU on an x86 system with one NVIDIA RTX PRO 6000 Blackwell Server Edition GPU, reach up to 131× faster inference compared to AlphaFold2. ARM-optimizations enable homology search on power efficient DGX Spark, and allow search beyond available GPU RAM on NVIDIA Grace Hopper Superchip at comparable overall folding speed. These accelerations enable high-throughput protein structure inference at no accuracy cost using familiar tools.

## Introduction

In 1972, Anfinsen proposed that protein structure is encoded in sequence [1]. Two decades later, Rost et al. achieved >70% secondary structure prediction using homology retrieval through multiple sequence alignment (MSA) generation and neural networks [2]. Morcos et al. later showed that statistical models leveraging MSAs could predict full 3D structures, including non-globular proteins [3], [4]. In 2021, Jumper et al. integrated massive data, compute, and deep learning (DL) to reach near-experimental accuracy with AlphaFold2 [5].

AlphaFold2’s success followed decades of expanding databases [6], faster MSA generation [7], and scalable DL [8], all powered by Moore’s law [9]. However, while databases grow exponentially [10] and new MSA and DL methods emerge [11], [12], Moore’s law may not hold forever [13].

Meeting future demand in protein structure prediction will require tighter hardware–software codesign to optimize throughput at inference. Current pipelines rely on two stages, i.e. MSA generation and transformer-based structure prediction, for instance in their baseline AlphaFold2 incarnation using MSA tools like JackHMMer [14] and HHblits [15], and DL inference through JAX.

Here, we focus on accelerations for deep learning-based inference using OpenFold [16] and TensorRT delivering 2.54×speedup compared to vanilla OpenFold, or 20.69× and 6.13×speedup compared to JAX-based AlphaFold2 and ColabFold-batch, respectively.

We further optimize MMseqs2-GPU [11] for execution on Blackwell and ARM, and interrogate end-to-end structure prediction on different hardware platforms. We find that end-to-end, OpenFold with TRT and MMseqs2-GPU on an x86-based server with one NVIDIA RTX PRO 6000 Blackwell Server Edition delivers the speediest protein folding pipeline achieving up to 131.4×speedup over the baseline AlphaFold2 pipeline, and 5.94× over the ColabFold [17] pipeline using MMseqs2-CPU. ARM-specific kernels enable larger-scale MSA generation on cohesive memory systems like the NVIDIA Grace-Hopper Superchip compared to x86+GPU-based systems, as well as efficient execution on small-form-factor systems like DGX Spark.

## Results

Following recent work [11] we focused on interrogating inference speed for protein structure prediction on 20 hard targets from CASP14 [18]. Protein structure prediction can be roughly broken up into two workloads: homology retrieval via the generation of Multiple Sequence Alignments (MSAs), representing the data pre-processing step, and Deep Learning (DL) inference, predicting structures from generated MSAs.

Metagenomic-scale GPU-based homology retrieval was enabled in *ColabFold-search* through MMseqs2-GPU via the introduction of GPU kernels for ungapped and gapped alignment [11]. These kernels were shown to reach lowest MSA generation time on an x86-based system equipped with an NVIDIA L40S GPU. We introduce NVIDIA Blackwell optimizations in MMseqs2-GPU enabling 1.4× faster MSA generation on an x86 system with one RTX PRO 6000 compared to x86+L40S. Compared to prior methods, this implementation is 191.4× faster than x86-based JackHMMER+HHblits used in the baseline AlphaFold2 pipeline, and 5.8× faster than ColabFold-search leveraging MMseqs2-CPU (Fig. 1).

**Fig. 1.**
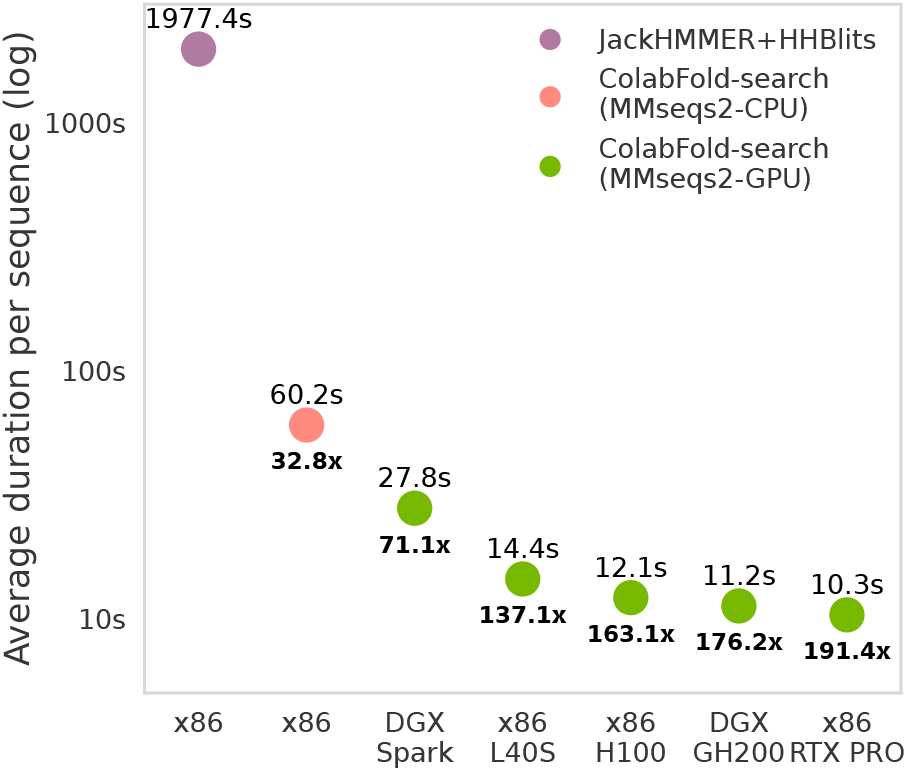
RTX PRO 6000 delivers fast MSA generation. An x86 server equipped with one RTX PRO 6000 delivers 1.4× faster metagenomic homology search with MMseqs2-GPU [11] compared to the previous best L40S. Grace-Hopper (GH200) reaches comparable speed with MMseqs2-GPU thanks to ARM-compilation improvements and high-bandwidth chip-to-chip (C2C) CPU-GPU interconnect.

Furthermore, we improved CPU cycle efficiency of MMseqs2-GPU on ARM by enabling 256-bit SIMD vector compilation. This allowed execution of structure prediction end-to-end on small-form factor systems like DGX Spark, and allowed Grace-Hopper (GH200) to reach a 1.29× speedup compared to x86+L40S (Fig. 1). MMseqs2-GPU on GH200 and DGX Spark takes advantage of unified memory and fast chip-to-chip interconnect alleviating GPU memory constraints. Using a representative benchmark set of 6370 queries against a reference set of 30M sequences, an x86 system with one NVIDIA L40S will degrade in throughput once the addressable space exceeds GPU memory of 48GB. Conversely, GH200 shows consistent throughput beyond GH200 GPU capacity of 96GBs (Fig. 2).

**Fig. 2.**
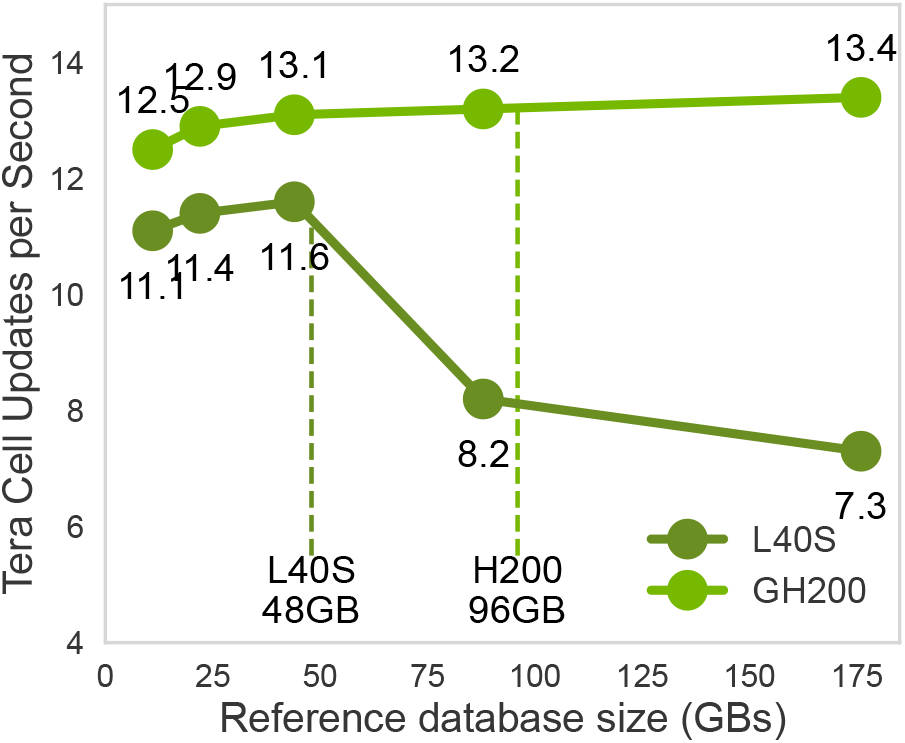
Grace-Hopper resilient to database scaling beyond GPU memory. Homology retrieval with MMseqs2-GPU on x86 systems is limited by available GPU memory, e.g. on an x86 system with a single L40S performance measured in Tera Cell Updates Per Second (TCUPS) drops 1.68x when exceeding available GPU memory (48GBs). This can be alleviated in multi-GPU systems by aggregating memory for all GPUs, and on ARM-based Grace-Hopper (GH200) through CPU-GPU shared memory and fast chip-to-chip (C2C) interconnect. GH200 maintains execution speed beyond available GPU memory (96GBs) using the host memory as extension.

Our primary contributions focus on protein folding DL inference. We optimised OpenFold [16], an open source re-implementation of AlphaFold2, offering a PyTorch-based backend with the ability to load pretrained AlphaFold2 weights. Starting from baseline OpenFold-PyTorch, we applied increasing levels of optimizations and evaluated inference speed using MSAs generated by the ColabFold-search pipeline leveraging MMseqs2-GPU for the 20 CASP14 targets. We monitored consistency in prediction accuracy from optimizations via TM-scores of the predicted structures against the ground truth CASP14 PDBs [19]. OpenFold compiled with TensorRT at BF16 precision (*OpenFold-TRT*) resulted the fastest AlphaFold2-style inference software, overall fastest on GH200 requiring on average 5.4s to fold proteins from their MSAs (Tab. I). NVIDIA RTX Pro 6000 closely followed requiring on average 5.6s (Fig. 3 & Tab. I). On these two systems, DL inference with OpenFold-TRT was 2.51× faster than vanilla OpenFold-PyTorch (Tab. I). Compared to baselines, on an RTX PRO 6000, OpenFold-TRT was 6.13× and 20.69× faster than ColabFold-batch and AlphaFold2, respectively (Fig. 3).

**TABLE 1.**
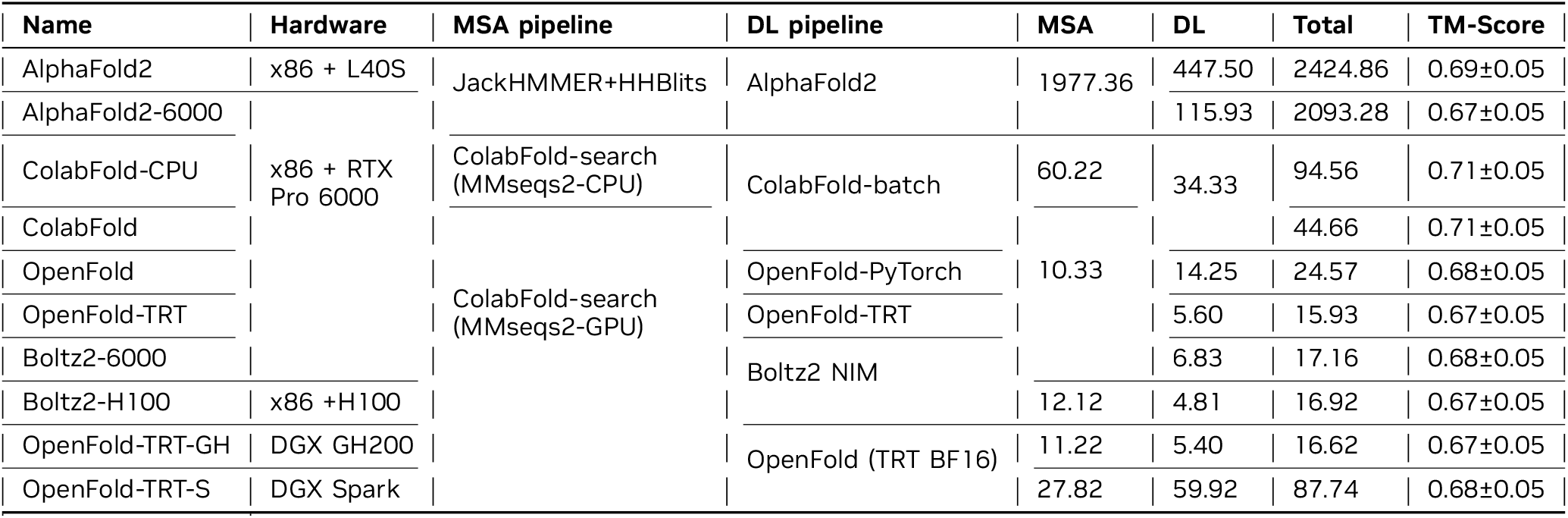
OpenFold-TRT improves DL inference across hardware configurations and delivers speediest end-to-end protein structure prediction. Average execution speed (s) for protein folding through MSA generation tool, DL inference pipeline and hardware configurations. Among AlphaFold2-like DL inference solutions, *OpenFold-TRT* reachest the best performance (column “Total”, bold). Boltz2 NIM reaches the overall fastest DL inference using H100 (column “DL”, bold), but loses end-to-end due to slower MSA generation time compared to RTX PRO 6000. All methods perform equally accurate.

**Fig. 3.**
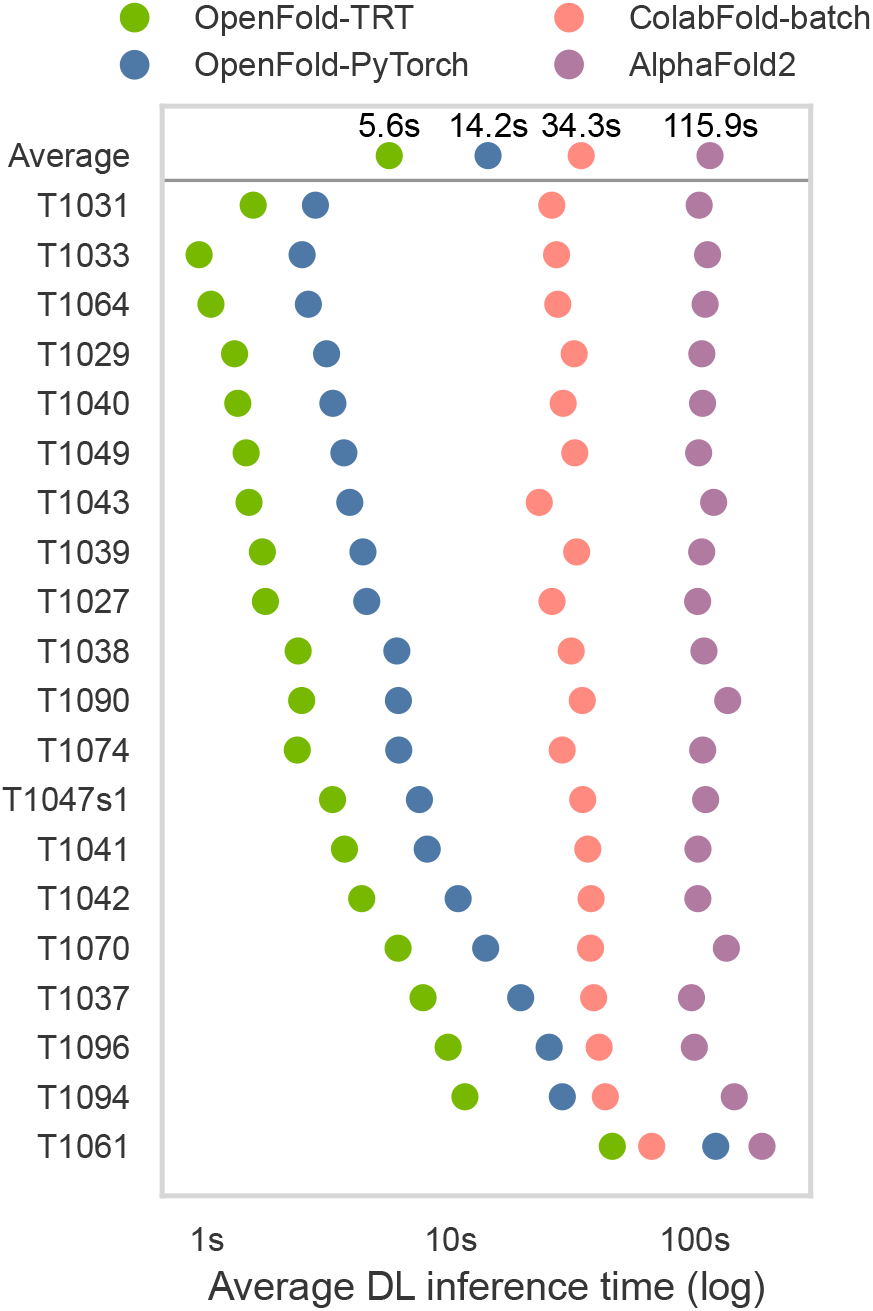
OpenFold-TRT reaches fastest DL inference. Consistently, OpenFold-TRT reaches the lowest inference speed across 20 CASP14 targets dependent on sequence length (targets ordered by sequence length shortest on top to longest at the bottom). On average, OpenFold-TRT DL inference is 20.69× faster than Alphafold2, and 2.54× faster than OpenFold-PyTorch.

One constraint when using TensorRT is its requirement to pre-allocate GPU memory; this allows for efficient inference, but conversely limits the maximum sequence length that can be inferred on. For instance, OpenFold-TRT on an L40S with 48GBs of GPU memory ran out of memory when inferring the second longest target (496 residues), while H100 with 80GBs ran out of memory when inferring the longest target (949 residues; Supplementary Data). This issue can be alleviated through larger GPU memory, e.g. RTX PRO 6000 with 96GB was able to infer all 20 targets.

End-to-end, i.e. combining MSA generation and DL inference, an x86 system equipped with an RTX PRO 6000 using ColabFold-search with the MMseqs2-GPU backend for MSA generation and OpenFold-TRT for DL inference resulted fastest, on average taking 15.93s per inference for the 20 CASP14 targets (Tab. I; *OpenFold-TRT*). Using the same software stack (ColabFold-search & OpenFold-TRT), GH200 is faster for DL inference, but slower for MSA generation, closely following at 0.96× the speed of RTX PRO 6000 (Tab. I; *OpenFold-TRT-GH*). Faster DL inference is expected on GH200 with 1979 TFLOPS [20] BF16 performance compared to RTX PRO 6000’s peak 503 TFLOPS [21]. *OpenFold-TRT* results 131.4×, 5.9× and 2.8× faster than baselines *AlphaFold2-6000, ColabFold-CPU*, and *ColabFold* on the same x86 system with one RTX 6000 Pro Server, respectively (Tab. I). While TM-score is consistent between OpenFold experiments across platforms, we measured lower TM-score for one target (T1064) in OpenFold compared to ColabFold-batch and AlphaFold2 (Supplementary Data). This may be attributable to random initialization or weight choice as we compared running the first weight set for each DL inference solution rather than selecting the best result from the five available weight sets for each method.

Recently, AlphaFold3 [22] was introduced promising faster DL inference through major architectural changes like dropping column-attention in the MSA representation, the use of PairFormer instead of Evoformer, and the removal of SE(3) equivariance from the structure module. Due to licensing considerations, we opted to evaluate Boltz-2 [12], an open source re-implementation of AlphaFold3, as additional baseline. We utilized the *Boltz-2 NIM (v1*.*1*.*0)*, which on top of AlphaFold3’s architectural efficiency integrates accelerated triangular attention and multiplication kernels from cuEquivariance, and is compiled for inference on TensorRT. DL inference through Boltz-2 on H100 resulted 1.17× faster than OpenFold-TRT on an RTX PRO 6000, and similarly to previous results, 1.42× faster on an H100 compared to on an RTX PRO 6000 (Tab. I). Including MSA generation through ColabFold-search using MMseqs2-GPU, Boltz-2 is 1.01× faster on H100 compared to RTX PRO 6000, but given slower MSA generation on H100, *OpenFold-TRT* is 1.06× faster overall.

We demonstrated that on average an x86 system with one RTX PRO 6000 using ColabFold-search leveraging the MMseqs2-GPU backend for MSA generation and OpenFold-TRT for DL inference results the fastest hardware and software combination to infer structures for 20 CASP14 targets (Tab. I). For AlphaFold2-like predictions, it is 2.8×faster than the previous fastest ColabFold-search with MMseqs2-GPU and ColabFold-batch on the same GPU. Considering the AlphaFold protein structure database [23] attempted to predict 350M sequences, at 44.66 seconds per inference using the previous fastest solution, this would require approximately 500 years to complete on one server with one GPU. Using multiple servers totaling 500 GPUs, it may take one year. Using the same infrastructure and OpenFold-TRT would require four and a half months, enabling faster large-scale in-silico data generation crucial to new generative methods for protein design [24], [25].

## Availability and Implementation

MMseqs2 and OpenFold are open source available software available at https://github.com/soedinglab/MMseqs2 and https://github.com/aqlaboratory/openfold, respectively. TensorRT is available at https://github.com/NVIDIA/TensorRT. Accelerations introduced in this work were upstreamed to MMseqs2, OpenFold and TensorRT, enabling public reproduction of these results provided system requirements are satisfied (CUDA 13.0 and TRT 10.13). Instructions for MMseqs2-GPU usage can be found at https://github.com/soedinglab/mmseqs2/wiki#gpu-accelerated-search, and instructions for using OpenFold-TRT can be found at https://openfold.readthedocs.io/en/latest/Inference.html#speeding-up-inference-with-tensorrt.

## Acknowledgments

We thank many NVIDIA colleagues for support, in particular Kyle Tretina, Kristopher Kersten, Milan Diebel, Varun Nanda Kumar, Neel Patel, Nick Venanzi, Kyle Gion, Duc Tran, and Scott Denton. M.S. acknowledges support by the National Research Foundation of Korea grants (2020M3A9G7103933, RS-2021-NR061659 and RS-2021-NR056571, RS-2024-00396026), Samsung DS Research Fund, Creative-Pioneering Researchers Program and Novo Nordisk Foundation (NNF24SA0092560). M.M. acknowledges support from the National Research Foundation of Korea (grant RS-2023-00250470). B.S. acknowledges support from the Deutsche Forschungsgemeinschaft (DFG, German Research Foundation) – project number 439669440 TRR319 RMaP TP C01. The funding body did not participate in the design of the study and collection, analysis, and interpretation of data and in writing the manuscript.

## Supporting Methods

### A. Deep Learning inference baselines

Protein structure prediction workflows can be divided into two principal stages: Multiple Sequence Alignment (MSA) generation and Deep Learning (DL) inference. To benchmark inference acceleration strategies independently of data preprocessing, we selected three established DL tools capable of loading the same set of AlphaFold2 weights. Notably, AlphaFold2 provides five distinct weight sets and produces five structures per input sequence; each structure can optionally be *relaxed* via short molecular dynamics (MD) simulations, with the relaxation step omitted in our study unless specified otherwise, since it is disabled by default in ColabFold-batch [17].

For comparability and reproducibility, we evaluated inference time and accuracy by running one protein sequence at a time using the *first* AlphaFold2 weight set in each tool, generating a single unrelaxed structure per query. This design ensures that reported performance metrics are independent of weight set sampling and relaxation overhead.

We benchmarked the following tools and configurations:

- **AlphaFold2 (JAX TF32)**: The canonical AlphaFold2 implementation in JAX, executed at TF32 precision using only the first weight set for each sequence.
- **ColabFold-batch (JAX BF16)**: ColabFold’s JAX variant with BF16 precision, leveraging the first AlphaFold2 weight set per sequence.
- **OpenFold (PyTorch TF32)**: The PyTorch-based OpenFold implementation, using the first AlphaFold2 weight set, with TF32 precision enforced.
- **OpenFold (TensorRT BF16)**: Our accelerated OpenFold compilation to TensorRT, inputting the first AlphaFold2 weight set, with the Evoformer core module executed at BF16 precision.

This framework allows systematic comparison of baseline AlphaFold2 and ColabFold-batch against PyTorch OpenFold and our TensorRT-optimized OpenFold pipeline, isolating the contribution of hardware and software accelerations to overall inference throughput and accuracy.

### B. OpenFold Inference Acceleration

The Evoformer block underpins the computational core of protein structure inference tools such as AlphaFold2 and OpenFold by applying advanced attention mechanisms to residue-residue interactions [16], [5]. To accelerate inference, we leveraged TensorRT optimizations focused on both precision management and dynamic handling of model inputs using OpenFold.

By default, OpenFold leverages PyTorch FP32 precision, with selective promotion to TF32 via `torch.set_float32_matmul_precision(`“`high`”) for improved computational throughput. Our TensorRT-compiled pipeline operates at mixed precision: TF32 for ExtraMSA and BF16 for the Evoformer module, which empirically achieves faster inference while maintaining prediction accuracy. To isolate the effect of hardware versus numeric casting, we ablated PyTorch’s explicit BF16 mode; although this provided some acceleration, the TensorRT pipeline yielded superior performance (see Supplementary Data).

To compile OpenFold with TensorRT, we utilize PyTorch’s TorchDynamo engine for ONNX export, which captures the model’s computational graph through bytecode analysis rather than traditional tracing. This approach preserves the dynamic nature of the Evoformer’s attention mechanisms while enabling subsequent TensorRT optimization. The ONNX export process leverages TorchDynamo with custom dynamic shape specifications for the Evoformer inputs:

~~~
evoformer_dynamic_shapes = { “m”: {1: seq_dim},
  % MSA representation
  “z”: {0: seq_dim, 1: seq_dim},
  % Pairwise representation “msa_mask”: {1: seq_dim},
  % MSA attention mask
  “pair_mask”: {0: seq_dim, 1: seq_dim}
  % Pair attention mask
}
~~~

For flexible execution, sequence length bounds are specified from S_MIN = 16 up to S_MAX (typi-cally set as `config.trt.max_sequence_len`), and optimization profiles are generated to efficiently cover single (i.e., [S_MIN, S_MAX]), dual (i.e., [S_MIN, S_MAX/2] and [S_MAX/2+1, S_MAX]), or four-range (i.e., four distinct ranges optimizing for different sequence length distributions) dynamic shapes. At inference time, model input tensors are matched to the appropriate profile automatically; out-of-profile queries either trigger a strict error (i.e., `ShapeError`), or fallback to PyTorch execution if enabled, ensuring robustness across diverse sequence lengths. Feature channels and MSA depths remain fixed (256 for MSA, 128 for pairs, 516 Evoformer stack, 5120 extra MSA stack). Further acceleration is achieved through TensorRT’s kernel fusion, which consolidates multi-step attention operations into optimized GPU kernels for reduced memory traffic and improved arithmetic intensity [26]. Persistent engine caching ensures rapid and reproducible deployment.

In essence, our proposed accelerations resolve to:

- **Precision setting:** using mixed-precision inference, i.e. TF32 for ExtraMSA, and BF16 for Evoformer
- **Dynamic Shape Support:** using dynamic profiles for efficient handling of variable-length protein sequences without requiring recompilation
- **Kernel Fusion:** multi-op attention blocks are fused into single GPU kernels to reduce bottlenecks and maximize performance

### C. MMseqs2-GPU improvements

GPU-accelerated homology search through MM-seqs2 was recently introduced via gapped and ungapped Gotoh-Smith-Waterman alignment kernels [27]. Here, we focused on improving these kernels for execution on Blackwell GPUs leveraging the introduced a new class of dynamic programming instructions (DPX) instruction set, and improving the CPU code generation of MMseqs2 on ARM-based systems:

- **Gapless alignment on Blackwell:** MMseqs2-GPU [27] used a novel implementation of a gapless pre-filter on GPUs to accelerate the protein database search simplifying the dynamic programming (DP) dependency scheme to only the diagonal neighbor. This allowed row-by-row processing of the DP matrix with each row partitioned among CUDA threads that are responsible for up to 128 cells each. Blackwell-based RTX PRO 6000 Server introduced a new class of dynamic programming instructions (DPX) instructions [28] and increased ALU pipes doubling the integer operation throughput compared to Ada-based L40S. To achieve peak performance, optimal distribution of the number of DP cells per CUDA thread is crucial. We analyzed the distribution of DP tile sizes and register usage per thread specifically for the Blackwell architecture and select the optimal to maximize alignment throughput. We thus employed 16-bit integers, where each DP-value is packed into a 32-bit word using s16x2 data types. The wider representation range of the DP-values allows us to search longer sequences than previous Ada generation, reaching higher peak performance.
- **MMseqs2 ARM optimizations:** To optimize MM-seqs2 execution on ARM-based architectures such as NVIDIA Grace-Hopper Superchip, we adapted core sequence alignment kernels and improved vectorization in expensive computational loops. To minimize execution overhead, rather than directly mapping x86 SSE to ARM NEON instructions, we re-implemented SIMD operations of `compare+all`, `any` and `horizontal max` using native NEON instructions (e.g., leveraging `UMINV` and `UMAXV`, which do not have direct SSE equivalents). These modifications reduced instruction latency and shortened dependency chains, enabling faster loop execution and improved single-thread throughput for both gapped and ungapped alignment routines. Performance was further enhanced by introducing support for 256-bit vector operations throughout the alignment workflow, implemented with pairs of 128-bit ARM NEON in-structions enabled by SIMDe macros and modern compilers (Clang ≥20.1.5). This approach allows better CPU utilization, nearly doubling vector compute throughput in expensive computational loops on modern ARM CPUs with 4x-6x FP/ASIMD pipelines. Profiling on representative protein datasets confirmed that these changes brought ARM execution efficiency to parity with the best x86 SSE implementations, with application-wide improvements exceeding 65% over the baseline code.

### D. Homology search pipelines

For the purpose of generating MSAs at metagenomic database scale as input to deep learning inference, we compared three pipelines:

- **JackHMMER+HHBlits:** uses HHblits [15] and JackHMMER [14] to generate three MSAs by searching the following databases: UniRef90 2020_01 (JackHMMer; w/ 139M sequences), MGnify 2018_12 (JackHMMer; w/ 287M sequences), Uniclust30 2018_08 (HHblits; w/ 15M profiles and 124M sequences total), BFD first release (HHblits; w/ 66M profiles and 2.1B sequences total). This search approach, databases and tools are represent the baseline in Alphafold2, where the three MSAs are combined into one to infer structures.
- **ColabFold-search (MMseqs2-CPU):** uses MM-seqs2 with default *k*-mer filtering on CPU to generate MSAs by first searching UniRef30 2023_02 (29M representatives and 277M sequences total) and then expanding from seed sequences identified in UniRef30 to Colab-FoldDB 2021_08 (209M representatives and 739M sequences total).
- **ColabFold-search (MMseqs2-GPU):** uses MM-seqs2-GPU with gapless filtering on the GPU to generate MSAs by searching the same databases as ColabFold-search (MMseqs2-CPU), using the same cascaded approach.

### E. Target sequences

To assess speed of MSA generation, MSA consistency, speed of deep learning inference, Template Modelling (TM) score consistency, and overall TM-score against ground truth we used a commonly used dataset of 20 “hard” targets from the CASP14 [18] competition. These targets ranged from 95 to 949 residues.

### F. Database scaling

As protein sequence database growth outpaces Moore’s law, efficient software solutions like MM-seqs2-GPU [27], searching protein databases very quickly on GPUs, are needed. In MMseqs2-GPU, when databases and clustering tricks, e.g. as employed by ColabFold-search through cascaded searching, outgrow available GPU memory, users can resort to either multi-GPU inference to maintain speed, pooling all available GPU memory on a node to grow the effective available total space, or resort to swapping between GPU and CPU in x86-based systems at a computational efficiency cost. In particular, MMSeqs2-GPU provides a mechanism to pre-load databases on GPU memory to prevent overhead for CPU-GPU data transfers across queries. In the scenario where the reference database is larger than device memory, the CPU-GPU bandwidth becomes the rate limiter for end-to-end performance, due to the requirement to stream the complete reference set in blocks from host to device memory to complete each query analysis.

Through the MMseqs2-GPU ARM optimizations introduced earlier we enabled ARM-based systems like the Grace Hopper Superchip, which fuses CPU and GPU with high bandwidth chip-to-chip (C2C) interconnect at 450 GB/s. This effectively alleviates this rate limiter otherwise observed on x86-based systems.

To analyze this behavior, we conducted a evaluation of the performance impact on database scaling for two different hardware platforms:

- AMD 2x64 cores and a L40S with a PCIe gen5 interconnection at 32GB/s
- NVIDIA Grace-Hopper (GH200) with 72 ARM v9 cores and a NVIDIA Hopper H200 with high bandwidth chipt-to-chip (C2C) CPU-GPU direct interconnection at 450 GB/s

We leveraged a commonly used dataset of 6370 full-length queries from UniProt against a reference set of 30M sequences. We increased the size of the reference set by replicating the set from one up to 16 times, which resulted in disk utilization from 11GB to 176GB.

### G. Hardware configurations

We evaluated different hardware platforms for both the MSA generation and DL inference pipelines. We restricted our analyses to systems with a single GPU. We focused our discussion on the best performing platforms, and relegated further results in Supplementary Data. We leveraged the following hardware platforms:

- **x86+L40S:** 1xNVIDIA L40S + AMD EPYC 7742 64-Core Processor (2 sockets total 128 cores) + 1TB RAM
- **x86+RTX PRO** 1xNVIDIA RTX PRO 6000 Black-well Server Edition + AMD EPYC 9554 64-Core Processor (2 sockets total 128 cores) + 2TB RAM
- **x86+H100** 1xNVIDIA H100 PCIe + AMD EPYC 7742 64-Core Processor (2 sockets total 128 cores) + 1TB RAM
- **DGX GH200:** NVIDIA Grace Hopper Superchip (GH200) with 72 ARM v9 cores with 480GB of LPDDR5X RAM and one NVIDIA Hopper H200 with 96GBs of HBM3
- **DGX Spark:** NVIDIA DGX Spark (GB10) with 20 ARM v9.2 CPU Cores (10 High-Performance Cortex-X925 and 10 Low-Power Cortex-A725) with 128GB of unified and coherent LPDDR5x RAM between ARM CPU and the NVIDIA Black-well GB20b GPU. (pre-production board and software stack)

